# Heterogeneous primary nucleation determines Aβ42 fibril structure

**DOI:** 10.64898/2026.10.30.763098

**Authors:** Emil Axell, Sara Linse, Dev Thacker

**Author notes:** These authors contributed equally.

## Abstract

The current developments in the field of amyloid fibril structure determination has revealed multiple structures for peptides of identical sequence. This may reflect variations in sample environment or similar nucleation barriers for multiple folds. Here we investigate the influence of aggregating surface and during fibril formation from monomeric amyloid β peptide (Aβ42). The fibril structures formed in identical solutions in glass vials / bottles and PEGylated polystyrene 96-well plates display very different monomer fold within the fibrils. This is a clear indication of extremely slow homogeneous primary nucleation. Instead, heterogeneous primary nucleation at vessel walls or at the three-line interface between air, water and vessel wall dictate the outcome in each solution, where the initially formed structure is propagated through elongation and secondary nucleation. The results further imply that the conversion towards the thermodynamically stable fibril structure one is slow for Aβ42.

## Introduction

Amyloid polymorphism, i.e. the co-existence of fibrils of different structures in the same sample, can be at equilibrium if the stabilities of the separate structures differ by one kT or less. The observation of amyloid polymorphism (1; 2; 3; 4; 5; 6; 7; 8; 9; 10; 11)may in the case of in-vivo fibrils be a consequence of variations in microenvironment. For a pure in-vitro system, an observed polymorphism more likely reflects the co-existence of kinetic products. In support of kinetic control, the level of polymorphism and its distribution in vitro was recently found to depend on total monomer concentration, with more initial polymorphism at higher monomer concentration, i.e. at higher aggregation rate (12).

A related phenomenon, the observation of fibrils of different structures in separate samples of the same protein or peptide, or in replicates of the same solution (13), suggests that these are kinetic products. Kinetic products will over time convert to the thermodynamic product, although this may take a very long time relative to typical experimental time frames. In the case of α-synuclein, αSyn, the rearrangement towards the most stable form with the lowest solubility is relatively fast and has indeed been shown to occur over a time scale of a week (13). Similarly, for islet amyloid polypeptide (IAPP, also called amylin) the evolution of fibril morphology distributions was observed over a time scale of a few weeks (14).

The stability of an amyloid fibril at a given temperature and solution condition can be quantified in terms of its solubility (i.e. the monomer concentration in equilibrium with the fibril (15; 16; 17; 18; 19), resistance to fragmentation (20), or may be extrapolated from data covering the temperature at the mid-point of its thermal denaturation equilibrium (21), or the denaturant concentration at the mid-point of its chemical denaturation equilibrium.

Amyloid formation is a nucleated polymerization reaction, which means that the barrier for nucleation is higher than that for elongation. This has been known for Aβ42 peptide since the early 1990s (22), and its aggregation mechanism includes at least three steps: primary nucleation from monomers only, elongation of fibrils by monomer addition, and secondary nucleation from monomers in the presence of fibrils (23). The free energy barriers for the separate steps can be derived from concentration-dependent studies at multiple temperatures. For Aβ42 in 20 mM sodium phosphate, pH 8.0 in PEGylated polystyrene plates, this approach revealed the following estimates: primary nucleation 65 kJ/mol, secondary nucleation 15 kJ/-mol and elongation 5 kJ/mol (24). Nucleation in bulk solution has to our knowledge not been observed, whereas heterogeneous primary nucleation can be inferred from optical microcopy images (25). Thus, in the absence of any reports of homogeneous primary nucleation of Aβ42, the number for the primary nucleation barrier likely reflects heterogeneous primary nucleation at the PEG surface.

At least 6 structural models have been reported for Aβ42 fibrils formed fully *in vitro* as well as for brain-derived material. Inspection of these structures reveals a variation in the monomer fold and the number of monomers per plane. For example, the structure models based on solid-state NMR (ssNMR) for fibrils formed in neutral phosphate buffer display similar S conformation for a highly ordered core comprising residues 15-42, with few structural constraints for the 14 N-terminal residues suggesting the N-terminus is relatively unstructured (26; 27; 28). The finding of multiple identical ssNMR spectra and structural models for multiple independent samples across laboratories and within a laboratory suggests that this is a robust fold, either reflecting a thermodynamic minimum or that the same kinetic product is formed over and over again with similar buffer and surface. Using small-angle X-ray scattering, the structures of the cross-section with 4 monomers per plane could be modeled (29). Another S-shaped fold is observed for fibrils formed at acidic pH in 30 % acetonitrile (30), in which case all 42 residues seem to be ordered. A structural model for brain-derived fibrils has been obtained based on cryo-EM data (31) for which the structure of the N-terminus have been added by simulation(32) **Describe**. Which show more of a U conformation? In this work, the structures of fibrils formed from monomer in different containers but in identical or closely similar buffer conditions in terms of ionic strength and pH were compared using cryo-EM. We show that fibril structure and the monomer fold in the filament changes based on the container surface. Further, we perform seeding experiments to show that the structure adapted by the aggregating monomer must be compatible with the parent seed structure for efficient seeding to take place.

## 1 Methods

### Aβ42 expression and purification and monomer isolation

Aβ(M1-42), with the sequence MDAEFRHDSGYEVHHQKLVFFAEDVGSNKGAIIGLMVG-GVVIA, was expressed in *E. coli* BL21 DE3 pLysS star in LB medium with IPTG induction and purified using ion exchange and size exclusion chromatography as described (23; 33) and multiple identical monomer aliquots were lyophylized and stored at -20°C.

### Fibril formation

Aliquots from above were dissolved in 1 mL 6 M GuHCl and subjected to size exclusion chromatography (SEC) on a 10/300 mm Superdex75 column in each experimental buffer (20 mM sodium phosphate, 0.2 mM EDTA, pH 8.0, or 20 mM Tris, 40 mM NaCl,0.2 mM EDTA, pH 8.0. The concentration of Aβ42 was determined from the absorbance at 280 nm of the collected fraction, using ϵ280 = 1440 l mol^*−*1^ cm^*−*1^, after which the monomer was diluted to 5 µM in the same buffer as used in the respective SEC run. Fibril formation was initiated by placing the solution in wells of a PEGylated plate or in glass vials and incubation of these containers at 37°C under quiescent conditions.

### Cryo-EM sample preparation

Fibrils prepared in the various conditions as described above were used for sample preparation. The samples were plunged using an automated plunge freezer (Leica EM GP) set at 20 °C with 90% relative humidity. Thin liquid films were created on glow-discharged lacey carbon 300 mesh grids (Agar Scientific) and blotted with filter paper before being plunged into liquid ethane at -184°C, ensuring vitrification in a glass-like state that prevents ice crystal formation and preserves the original microstructures. The grids were stored in liquid nitrogen until further use.

### Cryo-EM data collection and processing

The cryoEM datasets were collected at the SciLife lab node in Umeå, Sweden, using a Titan Krios electron microscope (Thermo Fisher) operated at 300 kV with a Falcon4 detector and a Selectris energy filter (Thermo Fisher) was used operating with a 10 e^*−*^V slit. Datasets of approximately 10,000 movies were collected for all sample conditions, at a nominal magnification of 165,000x yielding a pixel size of 0.73 Å or 0.728 Å. A defocus range of -0.6 to -2.2 µm and a total dose of 40 e^*−*^/Å^2^ were used over an exposure time of *≈* 3 seconds.

TIFF compressed movies were motion corrected in RELION-5.1 (34) and CTF estimation was done for micrographs using CTFFIND4 (35). Autopicking of fibrils was performed using the amyloid picker integrated in RELION-5.1. Iterative rounds of 2D classification were performed to remove picking artifacts such as carbon edges and ice contamination, until only classes that could be protein fibrils remained. Polymorphs (if present) in the dataset were separated out and independent helical symmetry searches were performed for each polymorph using 3D classification. Electron density maps were generated using 3D auto-refine. Automated model building was performed using ModelAngelo, fixed using Coot, and refined using Phenix.

### Self- and cross-seeding

Fibrils prepared at each surface and solution condition were used to seed fresh monomers at the same or different condition, to estimate the balance between elongation and secondary nucleation on the one hand and de novo (primary heterogeneous) nucleation on the other.

In these experiments, the Aβ42 mnonomer concentration was 2.5 *µ*mM and the seed concentration was either 25 nM (1%) or 2.5 *µ*mM (50%). The samples were monitored using ThT fluorescence in PEGylated plates or quartz cuvettes, respectively, and the end state was imaged using cryo-TEM.

## 2 Results

The structure of fibrils formed under a set of conditions (Table 1) were investigated using cryo EM.

**Table 1:** Experimental conditions used for sample preparation.

| Sample | Buffer | Container / Surface material |
|---|---|---|
| 1 | phosphate | 96-well plates / PEGylate polystyrene |
| 2 | Tris | 96-well plates / PEGylate polystyrene |
| 3 | phosphate | borosilicate glass vial |
| 4 | Tris | borosilicate glass vial |
| 5 | phosphate | Protein LoBind tube / polypropylene |
| 6 | Tris | Protein LoBind tube / polypropylene |

### 2.1 Fibril morphology and structure determination by cryo-EM

Aβ42 aggregated to form fibrils under all condtions, as can be seen in Figure 1.

**Figure 1:**
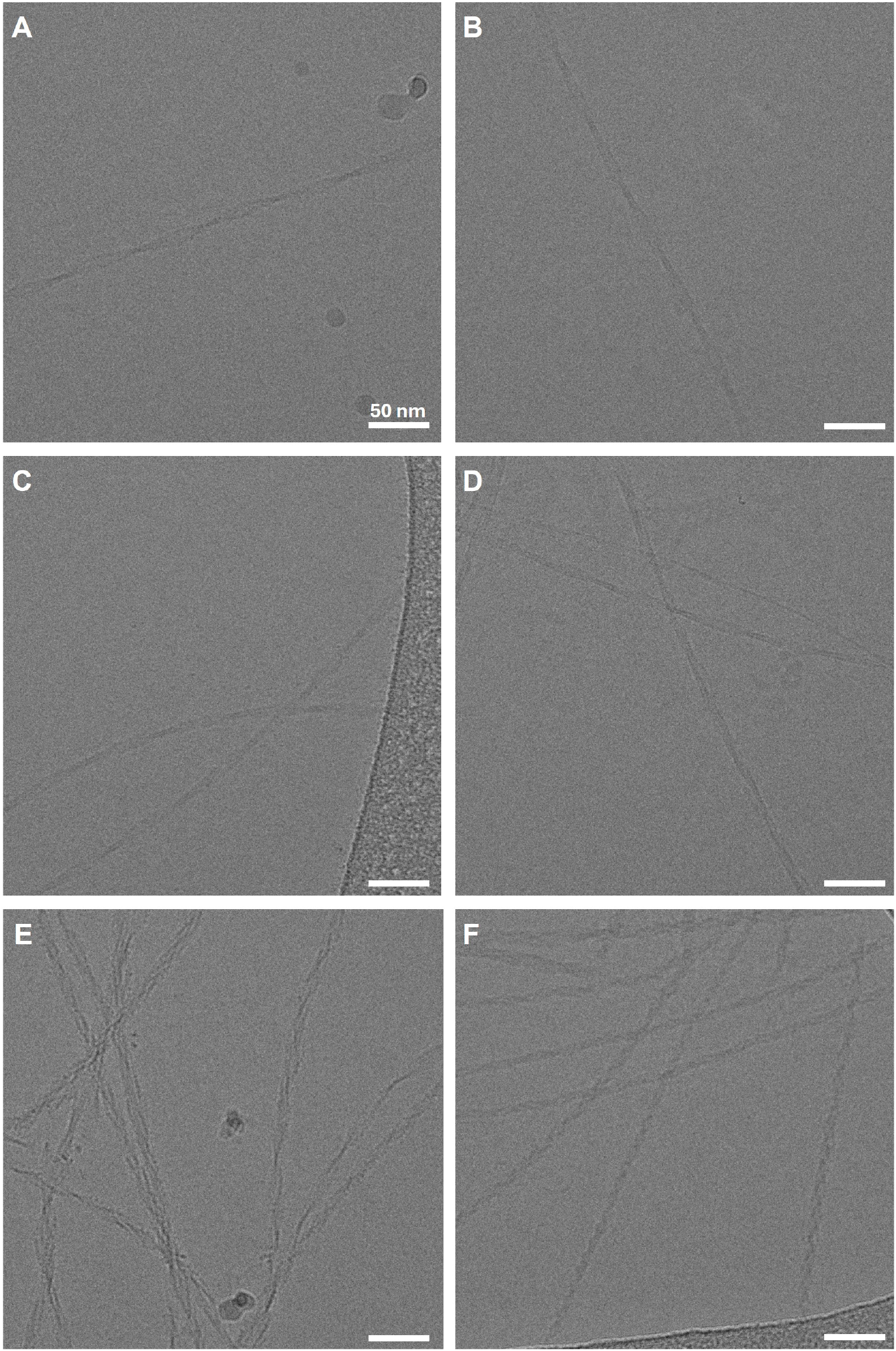
Cryo-EM images showing end-state fibrils assembled in **(A)** borosilicate glass vial in NaP buffer, **(B)** borosilicate glass vial in Tris buffer, **(C)** Protein LoBind tube in NaP buffer, **(D)** Protein LoBind tube in Tris buffer **(E)** 96-well plates in NaP buffer, and **(F)** 96-well plates in Tris buffer. The scale bar represents 50 nm.

To further investigate the structural characteristics of fibrils assembled under these conditions, we performed cryo-EM structure determination for the samples described above. The 2D classes are reported in Figure 2 for all six cases, i.e. Aβ42 fibrils formed in the presence of PEGylated polystyrene surfaces in NaP or Tris buffer, at identical ionic strength, pH 8.0, in the presence of glass surfaces in both buffers, as well as in Protein LoBind tubes in both buffers. 2D classes highlight the variations in the fibril morphologies across the six different sample preparation conditions, and show the polymorph population distribution in case of some of the samples. Fibrils assembled in plate readers display a polymorphic population, whereas to those in glass vials and LoBind tubes appear to be monomorphic.

**Figure 2:**
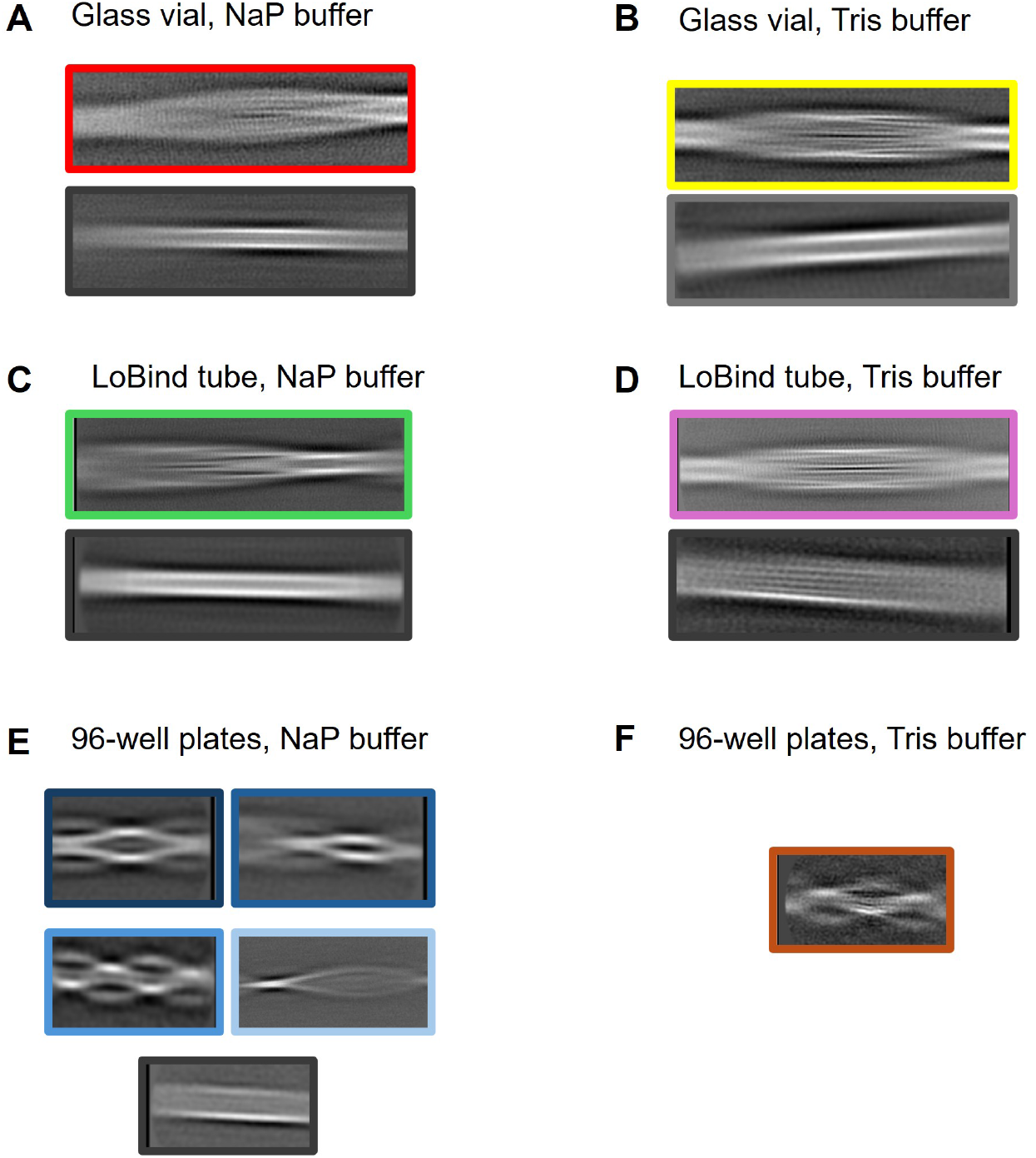
2D class averages from the processing of cryo-EM data highlight the difference in fibril morphology between different aggregating conditions. **(A)** borosilicate glass vial in NaP buffer, **(B)** borosilicate glass vial in Tris buffer, **(C)** Protein LoBind tube (polypropylene) in NaP buffer, **(D)** Protein LoBind tube in Tris buffer **(E)** 96-well plates (PEGylated polystyrene) in NaP buffer, and **(F)** 96-well plates in Tris buffer.

Once the polymorphs were separated based on their crossover distances from the 2D class averages, 3D classifications were performed to resolve their helical symmetry parameters. Figure 3 shows the resolved 3D classes from each dataset. Rermarkably, the fibrils assembled in 96-well plates show different monomer folds in both NaP and Tris buffers as compared to the fibrils assembled in borosilicate glass vials and LoBind tubes for both buffers. The fibrils from 96-well plates display a U-shaped monomer fold with different filament arrangements within each fibrils. For NaP buffer, we observe fibrils with 2, 3, and 4 monomers per plane of fibril. These monomers are ordered from Q15 to A42, with residues 1-14 forming the flexible fuzzy coat. In striking contrast, the fibrils observed in glass vial in Tris buffer, and LoBind tubes in NaP and Tris buffers consist of two completely ordered monomers per fibril plane, ie, residue 1 to 42 all forming part of the ordered fibril core. The reconstructed EM densities are shown in Figure 4 and the constructed models are shown in Figure 5. Even though they present with a different monomer fold, the 96-well fibrils and the LoBind tube fibrils have a structurally conserved monomer fold spanning 11 residues from A21 to I32, which are shown superimposed in Figure 5C. Interestingly, the fibrils formed in glass vials in NaP buffer have fewer ordered residues than those in glass vials in Tris buffer, but still a different monomer fold than the 96-well plate fibrils.

**Figure 3:**
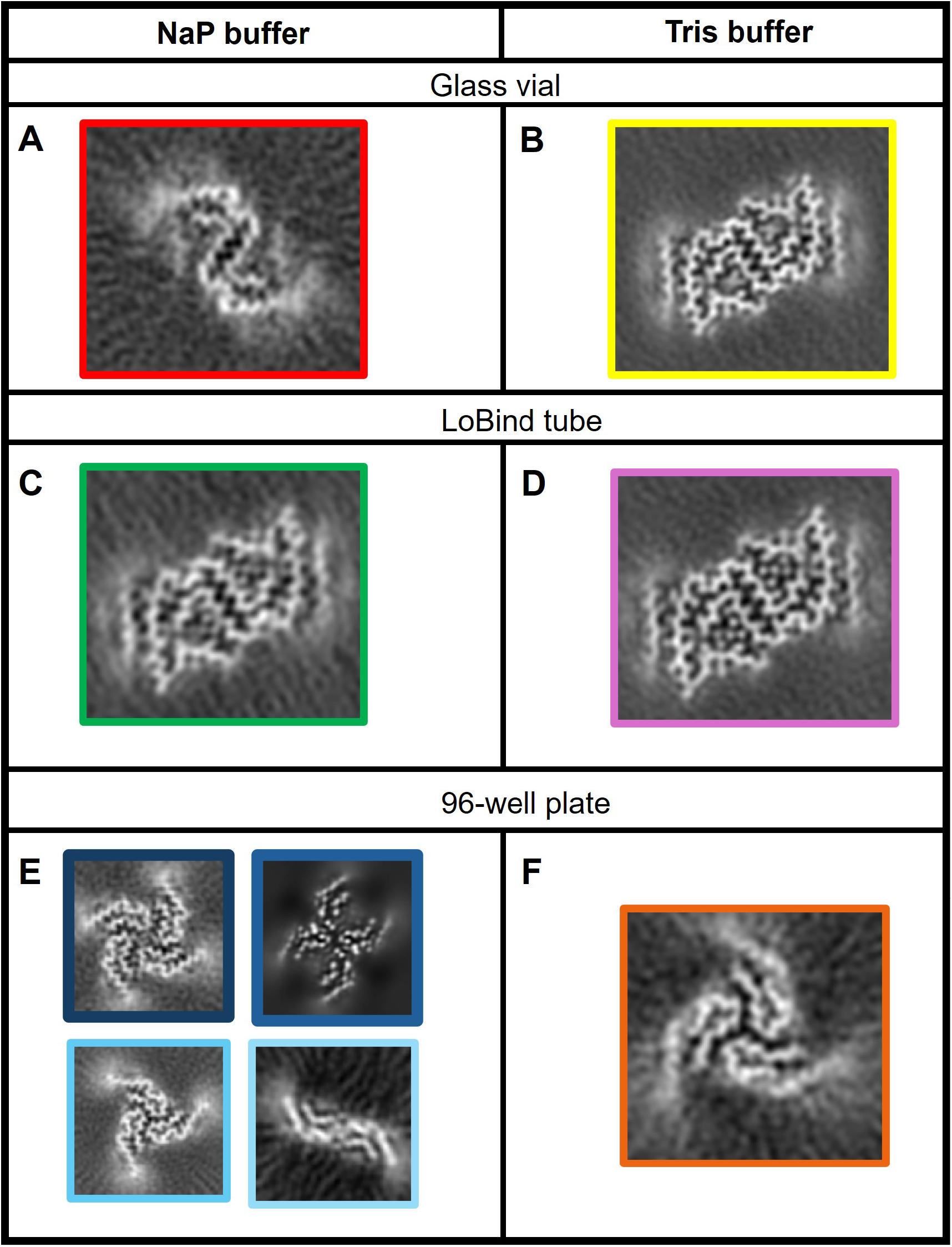
3D class averages showing the monomer folds within the fibrils assembled in each dataset across the different aggregation conditions. **(A)** borosilicate glass vial in NaP buffer, **(B)** borosilicate glass vial in Tris buffer, **(C)** Protein LoBind tube in NaP buffer, **(D)** Protein LoBind tube in Tris buffer **(E)** 96-well plates in NaP buffer, and **(F)** 96-well plates in Tris buffer.

**Figure 4:**
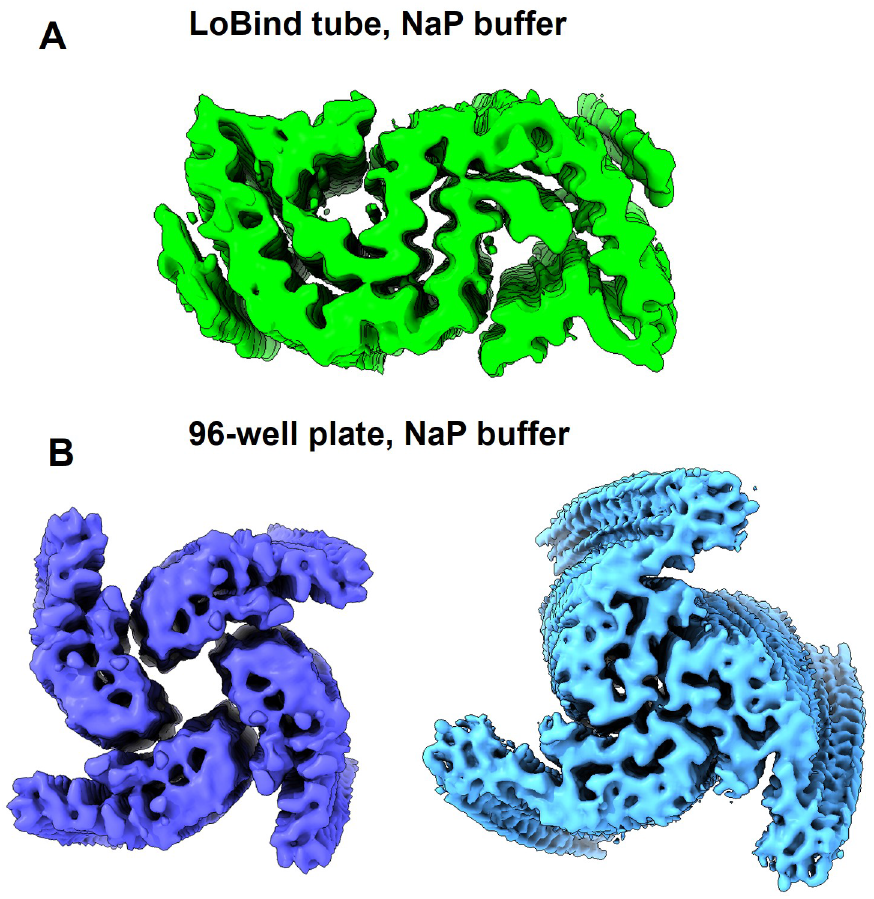
Reconstructed EM densities of fibrils assembled in **(A)** LoBind tubes in NaP buffer, and **(B)** 96-well plates in NaP buffer.

**Figure 5:**
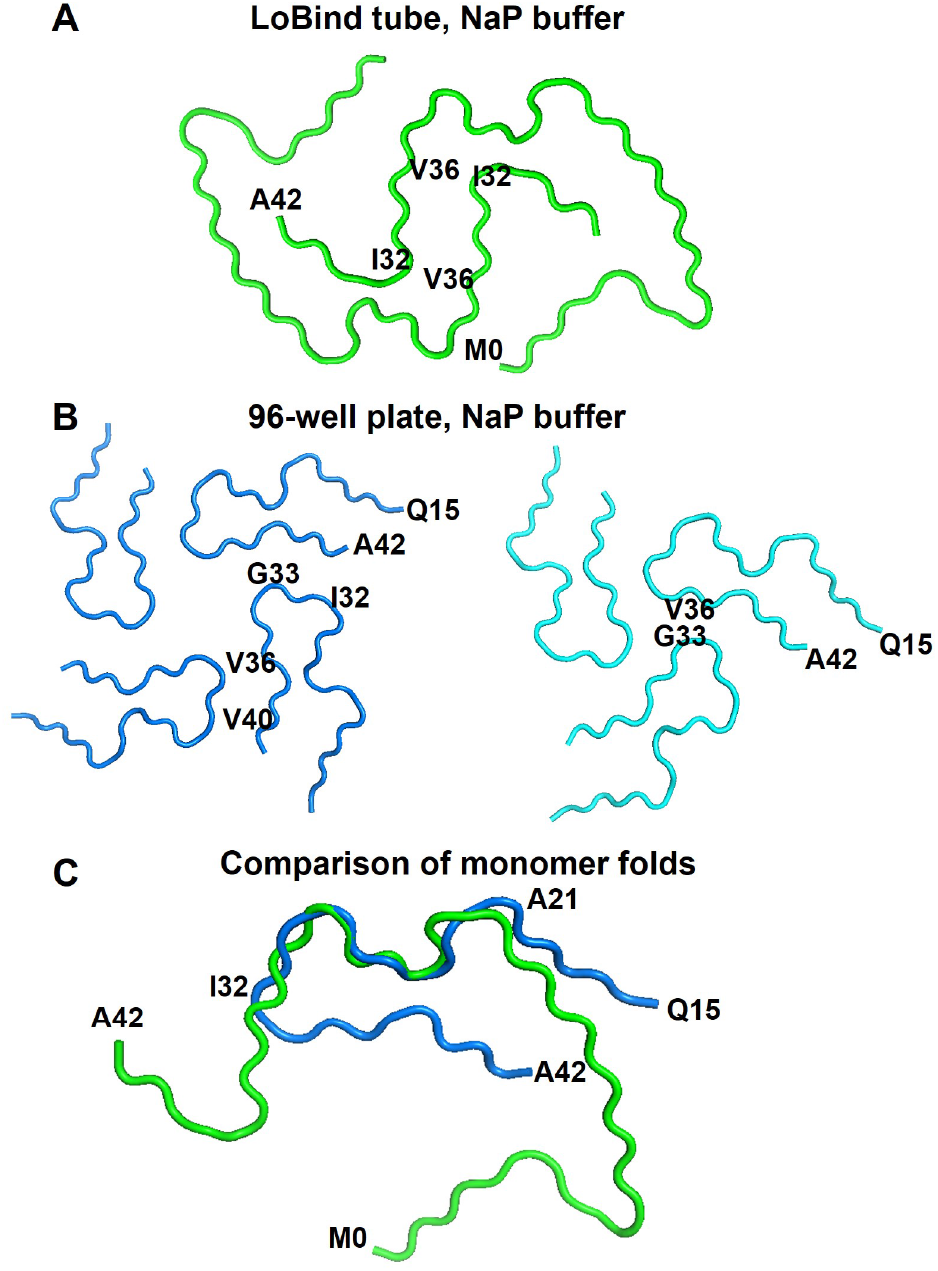
Models constructed for Aβ42 fibrils assembled in **(A)** LoBind tube in NaP buffer, and **(B)** two of the polymorphs observed in 96-well plates in NaP buffer. **(C)** Superimposed monomer folds of the above two conditions.

### 2.2 Cross-seeding with different morphologies

After having established that nucleation on different surfaces leads to distinct Aβ42 fibril folds, next we sought to investigate whether these fibrils also differed in their ability to seed new reactions. Fibrils formed under different solution and surface conditions were therefore used to seed freshly isolated Aβ42 monomers (Fig. 6A). To mitigate potential differences in the number of fibril ends arising from agitation (36), all fibril preparations used as seeds were generated under quiescent conditions. First, we used a low seed concentration of 1% monomer equivalents at a total Aβ42 concentration of 2.5 µM (Fig. 6B). In this regime, the reaction informs on the overall ability of the pre-formed fibrils to catalyze secondary processes such as secondary nucleation. Both fibrils formed in PEGylated polystyrene microplates in either NaP or Tris and borosilicate glass bottles accelerated aggregation relative to the unseeded control, while no clear difference was observed between the two buffer conditions, the seeds derived in plates accelerated the aggregation substantially more than the fibrils derived in glass, as reflected by the shorter half-times (Fig. 6B,D). In contrast, no apparent difference in seeding potency was observed between NaP and Tris within the same vessel type. Thus, under low seed regime, the surface on which the monomers underwent fibril formation appeared to have a bigger influence on their subsequent seeding potency than the buffer used during fibril formation.

**Figure 6:**
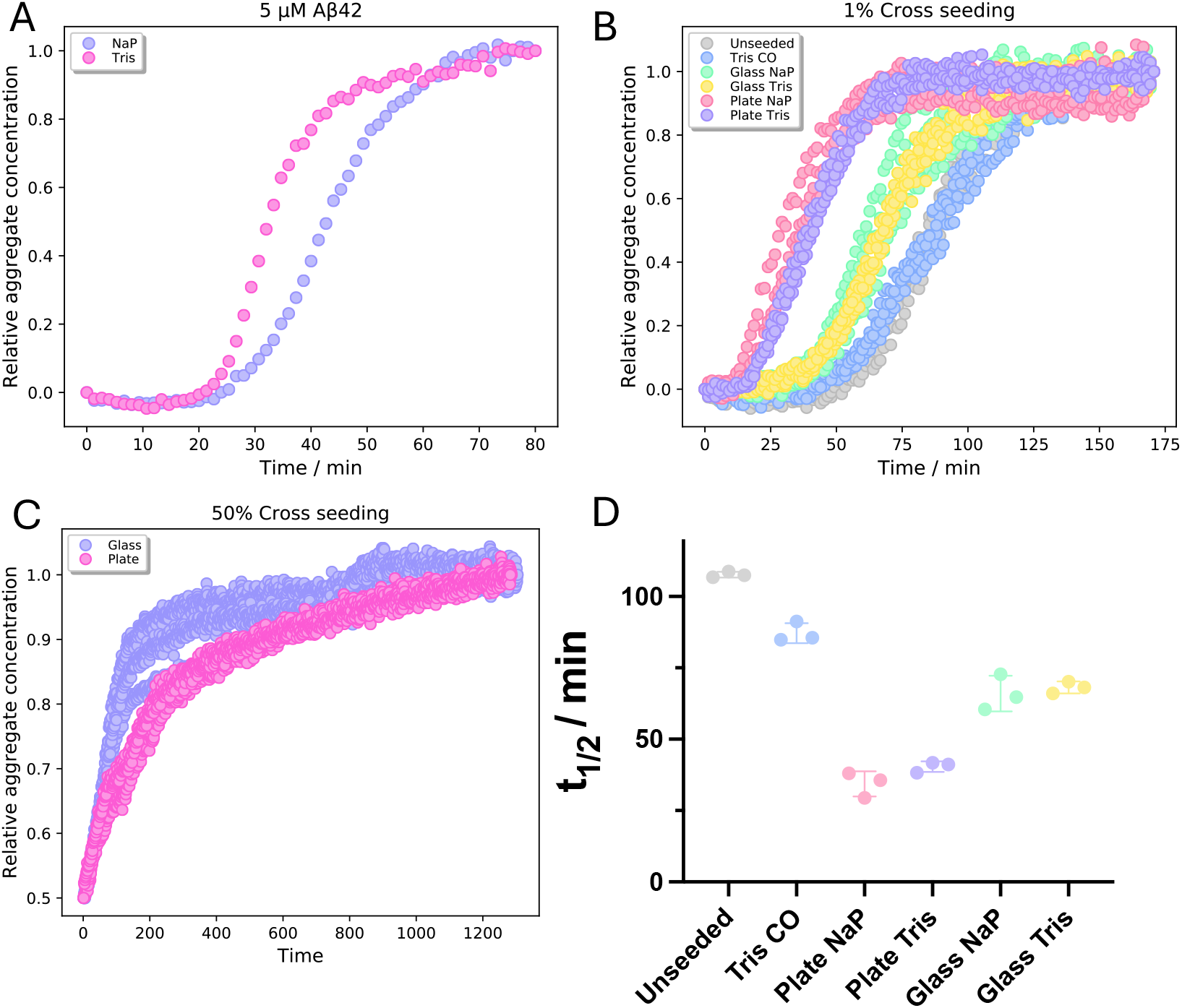
Polymorph-dependent seeding. (A) Seed formation: 5 µM Aβ42 monomer was incubated at 37 °C in either 20 mM sodium phosphate, 0.2 mM EDTA, pH 8.0 (NaP, purple), or 20 mM Tris, 40 mM NaCl, 0.2 mM EDTA, pH 8.0 (Tris, pink). Fibril formation was monitored with ThT fluorescence. The fibrils used for seeding in (B) were formed quiescently without ThT for 48 h and were made either in Corning 3881 microplates (plate) or in borosilicate glass bottles (glass). **(B) 1% cross-seeding:** Kinetic traces of freshly isolated Aβ42 monomers (2.5 µM total concentration) in NaP supplemented with 1% fibrils formed either in glass or plate and in either NaP or Tris. Unseeded Aβ42 in NaP (gray) and a Tris carryover control (blue), containing the same volume of Tris buffer introduced with the Tris seeds but without fibrils, were included as controls. Seeds formed in glass in NaP and Tris are shown in green and yellow, respectively, and seeds formed in microplates in NaP and Tris in pink and purple. **(C) 50% cross-seeding:** Freshly isolated Aβ42 monomers were supplemented with NaP fibrils formed either in microplates or glass to a total Aβ42 concentration of 2.5 µM and a seed fraction of 50%. Seeds formed in glass are shown in purple and seeds formed in microplates in pink. **(D) Half-times:** Half-times were determined from the normalized 1% seeding reactions in (B). Individual replicates are shown with mean *±* SD. Kinetic traces in (A–C) were normalized in AmyloFit.

In order to investigate whether this difference was also apparent under strongly seeded conditions. Reactions with seed fraction to 50% (Fig. 6C) was carried out. Both seed populations were formed in NaP, and the reseeding reactions were performed under identical NaP conditions in the same microplate format to not introduce artifacts from introduction of Tris/-NaCl. At a total Aβ42 concentration of 2.5 µM, the 50% seed fraction leaves 1.25 µM soluble monomer at the start of the reaction. Under this regime, elongation on pre-existing fibrils is expected to consume more monomer than primary and secondary nucleation and contribute more to the observed kinetics. Interestingly, glass derived seeds showed a steeper initial increase in fluorescence than plate derived seeds, opposite to their relative potency in the 1% seeding experiment. Both reactions showed a slow approach towards the final fluorescence plateau. The difference observed under strongly seeded conditions therefore suggests that the greater potency of plate derived fibrils at 1% seed cannot simply be reconciled with a greater elongation rate. However, the observed rate at high seed concentration will depend on both the intrinsic elongation rate and the number of elongation competent fibril ends. Thus, the current data can therefore not allow the difference to be assigned solely to *k*_+_.

Taken together, the low- and high-seed experiments suggest that fibrils formed in containers of different surface materials display distinct propagation properties. Plate derived fibrils showed greater overall seeding potency at low seed concentration, whereas glass derived fibrils displayed faster initial growth under strongly seeded conditions.

## 3 Discussion

We performed cryo-EM structure determination of Aβ42 fibrils assembled in six different conditions, across two different buffers and three different aggregating surfaces. The structures from 96-well plates present a different monomer fold as compared to the other aggregating conditions. The structures of Aβ42 fibrils formed in the presence PEGylated polystyrene surfaces (Figure 5) are distinct from the ssNMR structure for Aβ42 fibrils formed in glass bottle (28). The S-like fold of the monomer within the fibril plane as observed in aggregates formed in glass is clearly not observed in the Aβ42 fibrils formed in PEGylated plates. In This is a clear sign of slow homogeneous primary nucleation in bulk, with faster heterogeneous primary nucleation dictating the outcome.

## 4 Conclusion

The structures of Aβ42 fibrils formed in the presence of glass or PEGylated polystyrene surfaces are very different in terms of the monomer fold within the fibril plane. This is a clear sign of slow or negligible homogeneous primary nucleation in bulk, with faster heterogeneous primary nucleation dictating the outcome.

